# Mutagenesis and homology modeling reveal a predicted pocket of lysophosphatidylcholine acyltransferase 2 to catch Acyl-CoA

**DOI:** 10.1101/2020.10.31.363515

**Authors:** Fumie Hamano, Kazuaki Matoba, Tomomi Hashidate-Yoshida, Tomoyuki Suzuki, Kiyotake Miura, Daisuke Hishikawa, Takeshi Harayama, Koichi Yuki, Yoshihiro Kita, Nobuo N. Noda, Takao Shimizu, Hideo Shindou

## Abstract

Platelet-activating factor (PAF) is a potent proinflammatory phospholipid mediator that elicits various cellular functions and promotes several pathological conditions, including anaphylaxis and neuropathic pain. PAF is biosynthesized by two types of lyso-PAF acetyltransferases: lysophosphatidylcholine acyltransferase 1 (LPCAT1) and LPCAT2, which are constitutive and inducible forms of lyso-PAF acetyltransferase, respectively. Because LPCAT2 mainly produces PAF under inflammatory conditions, understanding the structure of LPCAT2 is important for developing specific drugs against PAF-related inflammatory diseases. Although the structure of LPCAT2 has not been determined, the crystal structure was reported for *Thermotoga maritima* PlsC, an enzyme in the same enzyme family as LPCAT2. In this study, we identified residues in mouse LPCAT2 essential for its enzymatic activity and a potential acyl-coenzyme A (CoA)-binding pocket, based on homology modeling of mouse LPCAT2 with PlsC. We also found that Ala115 of mouse LPCAT2 was important for acyl-CoA selectivity. In conclusion, these results predict the structure of mouse LPCAT2. Our findings have implications for the future development of new drugs against PAF-related diseases.

## Introduction

Platelet-activating factor (PAF, 1-*O*-alkyl-2-acetyl-*sn*-glycero-3-phosphocholine) is a potent pro-inflammatory lipid mediator that activates the PAF receptor, which is a G protein-coupled receptor. PAF elicits several biological effects, such as platelet activation, airway constriction, hypotension, systemic anaphylaxis, and neuropathic pain (1, 2). In response to extracellular stimuli, 1-*O*-alkyl-2-acyl-*sn*-glycero-3-phosphocholine is hydrolyzed by phospholipase A2 to produce polyunsaturated fatty acid and 1-*O*-alkyl-*sn*-glycero-3-phosphocholine (lyso-PAF). Under these conditions, PAF is rapidly biosynthesized from lyso-PAF by lyso-PAF acetyltransferase [EC: 2.3.1.67] (3–5).

Previously, we identified two types of lyso-PAF acetyltransferases, lysophosphatidylcholine acyltransferase 1 (LPCAT1) (6) and LPCAT2 (5). LPCAT1 is a constitutively expressed lyso-PAF acetyltransferase that also generates dipalmitoyl-phosphatidylcholine (PC) through its LPCAT activity [EC: 2.3.1.23] (6–8). Dipalmitoyl-PC is a main component of pulmonary surfactant, which is essential for alveolar gas exchange (9, 10). In contrast, LPCAT2 expression is regulated in three independent manners in macrophages. In a previous study, LPCAT2 was rapidly phosphorylated and activated by protein kinase C alpha following stimulation with PAF or ATP for 1 min (11). Lipopolysaccharide stimulation for 30 min and 16 h induced LPCAT2 phosphorylation by MAP kinase-activated protein kinase 2 and LPCAT2 expression (mRNA and protein), respectively, via Toll-like receptor 4 (5, 12). Although phosphorylation of residue Ser34 in LPCAT2 is common, the responsible kinase and the time course of such phosphorylation are different. Thus, LPCAT2 acts as an inducible and activated form of lyso-PAF acetyltransferase under inflammatory conditions. LPCAT2 also possesses LPCAT activity that catalyzes acyl-coenzyme A (CoA) incorporation into membrane phosphatidylcholine (PC). However, the biological roles of this LPCAT2 activity, which incorporates long fatty acids remain unknown, because LPCAT2 deficient mice showed reduction of PAF, but not other phospholipids (2). Whereas, it is reported that LPCAT2 produces PC at lipid droplet (13, 14).

Both LPCAT1 and LPCAT2 are members of 1-acylglycerol-3-phosphate *O*-acyltransferase (AGPAT) family. Several groups (including our group) have reported AGPAT motifs that are essential for acyltransferase activity (6, 15–17). To develop drugs for PAF-related diseases, it is important to understand the biochemical characteristics of PAF production and the structure of LPCAT2. However, the LPCAT2 structure is unknown because it is difficult to solubilize and purify LPCAT2 from the membrane fraction of tissues or cells. Recently, Robertson et al. reported the crystal structure of *Thermotoga maritima* PlsC, which is a member of the AGPAT family and possesses lysophosphatidic acid acyltransferase activity (18). This report is useful for analyzing the structure of LPCAT2.

In this study, site-directed mutagenesis revealed AGPAT motifs in mouse LPCAT2 (mLPCAT2) and important residues for its acyl-CoA selectivity. Additionally, utilizing the information from the PlsC crystal, an acyl-CoA binding pocket of mLPCAT2 was predicted by homology modeling. Our data should help in analyzing the biochemical characteristics of mLPCAT2 and in developing new drugs for PAF-related diseases.

## Results

### Alignment and homology modeling of mouse LPCAT1 (mLPCAT1), mLPCAT2, and *Thermotoga maritima* PlsC (tmPlsC)

tmPlsC (18) is homologous to mLPCAT1 and mLPCAT2, and its structure is available in the Protein Data Bank (PDB); thus, the tmPlsC structure was selected for predicting the structure of mLPCAT2. According to data from the EMBOSS Water website (https://www.ebi.ac.uk/Tools/psa/emboss_water/), tmPlsC shares 26% identity with both mLPCAT1 and mLPCAT2. The primary structure of mLPCAT2 was compared with those of mLPCAT1 and tmPlsC (Fig. 1A). AGPAT motifs 1 to 4 are highly conserved among all three enzymes. For homology modeling, we analyzed short forms of mLPCAT2 (residues 52–320) and mLPCAT1 (residues 44–301) because tmPlsC is shorter than both mLPCAT1 and mLPCAT2 (Fig. 1B). Consequently, the dilysine motif (which localizes proteins to the endoplasmic reticulum) and the EF-hand (like) motifs of mLPCAT1 and mLPCAT2 were not analyzed in this study, as tmPlsC does not possess these motifs (Fig. 1B). Phosphorylation of the N-terminal Ser34 residue of mLPCAT2 was also not analyzed during the homology modeling (Fig. 1A, B). AGPAT motifs 1–4, which have been identified by three independent groups (6, 15–17), are indicated in colored squares (green, magenta, blue, and orange). The red open squares show the amino acid residues analyzed by mutagenesis in this study (Fig. 1A). mLPCAT1 and mLPCAT2 showed high homology in both the AGPAT motif and the predicted loop region following motif 4 (red square in the loop). Homology modeling was performed using the tmPlsC structure (18) as a template (Fig. 1C, Supporting Information models 1, 2). These structures show that two N-terminal α-helices, which anchor tmPlsC to one leaflet of the membrane, were also predicted to be present in mLPCAT1 (residues 44–112) and mLPCAT2 (residues 52–124).

**Fig. 1.**
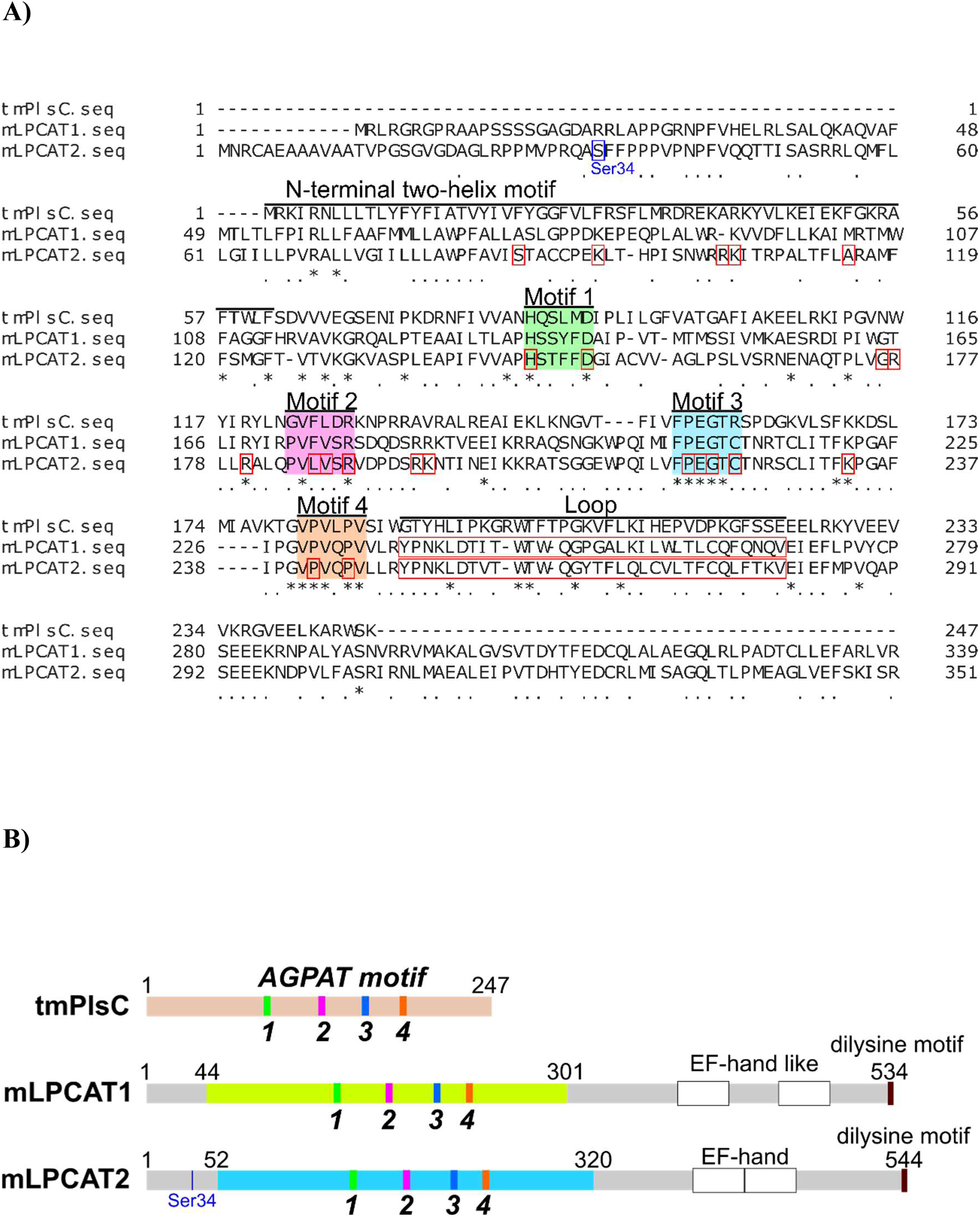
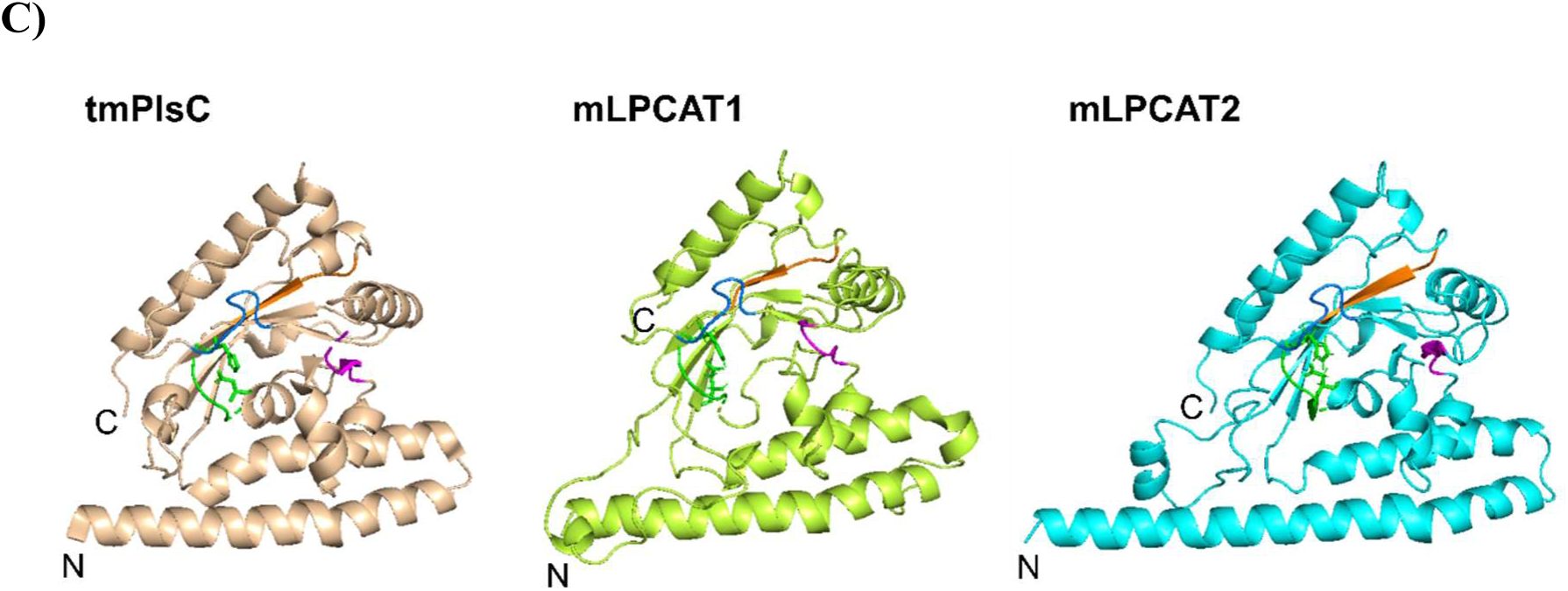
Alignment and homology modeling of tmPlsC, mLPCAT1, and 2 structures. A) A region of tmPlsC, mLPCAT and mLPCAT2 containing conserved regions, i.e., AGPAT motifs 1 (green), 2 (magenta), 3 (blue), and 4 (orange), was used as a query for GENETYX-MAC software. The amino acid residues analyzed by mutagenesis in this study are indicated with red open squares. Asterisks represent conserved residues, and dots represent weakly similar residues. B) Schematic representations of the structures of tmPlsC, mLPCAT1, and mLPCAT2. Motifs 1– 4 are colored green, magenta, blue, and orange, respectively. We modeled mLPCAT1 (light green domain) and mLPCAT2 (light blue domain) based on the crystal structure data for tmPlsC (beige). C) Homology modeling of mLPCAT1 and mLPCAT2. Ribbon models of tmPlsC (used as the template), mLPCAT1, and mLPCAT2. Motifs 1–4 are highlighted as follow: motif 1, green; motif 2, magenta; motif 3, blue; motif 4, orange.

### Identification of AGPAT motifs in mLPCAT2

Next, we evaluated the roles of the AGPAT motifs and the N-terminal two-helix motif. The following Ala or Glu substitutions were made to prepare mLPCAT2 mutants: H146A and D151A in AGPAT motif 1; G176A, R177A, R177E, R180A, L186A, V187A, R189A, and RK195/196AA around AGPAT motif 2; P219A, E220A, G221A, and C223A in AGPAT motif 3; K233A, P242A, and P245A around AGPAT motif 4; and K94A and RK104/105AA in the N-terminal two-helix motif (Fig. 1A and 2). The expression levels of all proteins used in these assays were almost similar to that of wild type (WT) mLPCAT2 (Supporting Fig. 1). Because mLPCAT2 possesses two types of activities, i.e., lyso-PAF acetyltransferase with acetyl-CoA and lyso-PAF acyltransferase with arachidonoyl-CoA, we assessed both activities of each mutant. Consistent with previous results for the AGPAT motifs (6, 15–17, 19), most mutations reduced both activities, confirming that these regions are essential for mLPCAT2 activities. Ala mutants of His146 and Asp151, which were located in the active center in motif 1, completely lost mLPCAT2 activity. Furthermore, the L186A, V187A, P219A, E220A, C223A, P242A, and P245A mutants also showed the loss of the activity. The R177A and R180A mutants retained half of the WT activity. According to our homology modeling for mLPCAT2 (Fig. 1C, 2A, Supporting Information model 2), Arg177 and Arg180 were located near the N-terminal two-helix motif. The positive charges of amino acids near the membrane may participate in interactions with the polar groups of membrane phospholipids and form contacts with the phosphate group of the acyl-CoA substrate. Thus, Arg177 was next substituted with the acid amino acid, Glu to examine whether the positive charge at this position is necessary for the activities of mLPCAT2. The R177E mutant lost mLPCAT2 activities, although the R177A mutant retained partial activities. These results suggest that the positive charge at this position largely contributes to the activities. Gly176 is somewhat distant from the active center; however, the G176A mutant also showed the loss of the activity. Because the Pro173– x174–x175–Gly176 sequence is highly conserved among various LPLATs and would provide a kinked structure between motif 1 and motif 2, the kink might be necessary for mLPCAT2 activities (red shading in Supporting Fig. 2). Each hydrophobic pocket was observed in our homology models (Fig. 3A, Supporting Information model 2). Arg187, Lys104, Arg105, and Lys223 were located on the upper enzyme surface, around the opening of the hydrophobic pocket (Fig. 3B, Supporting Information model 2), and these mutants lost almost all activity. However, the activities of the K94A and RK104/105AA mutants were partially retained. Residues Lys94, Arg104, and Lys105 were located in the N-terminal two-helix motif. Gly221 was near the active center; however, the G221A mutant retained approximately half of its activity, probably because the change from Gly to Ala was not significant.

**Fig. 2.**
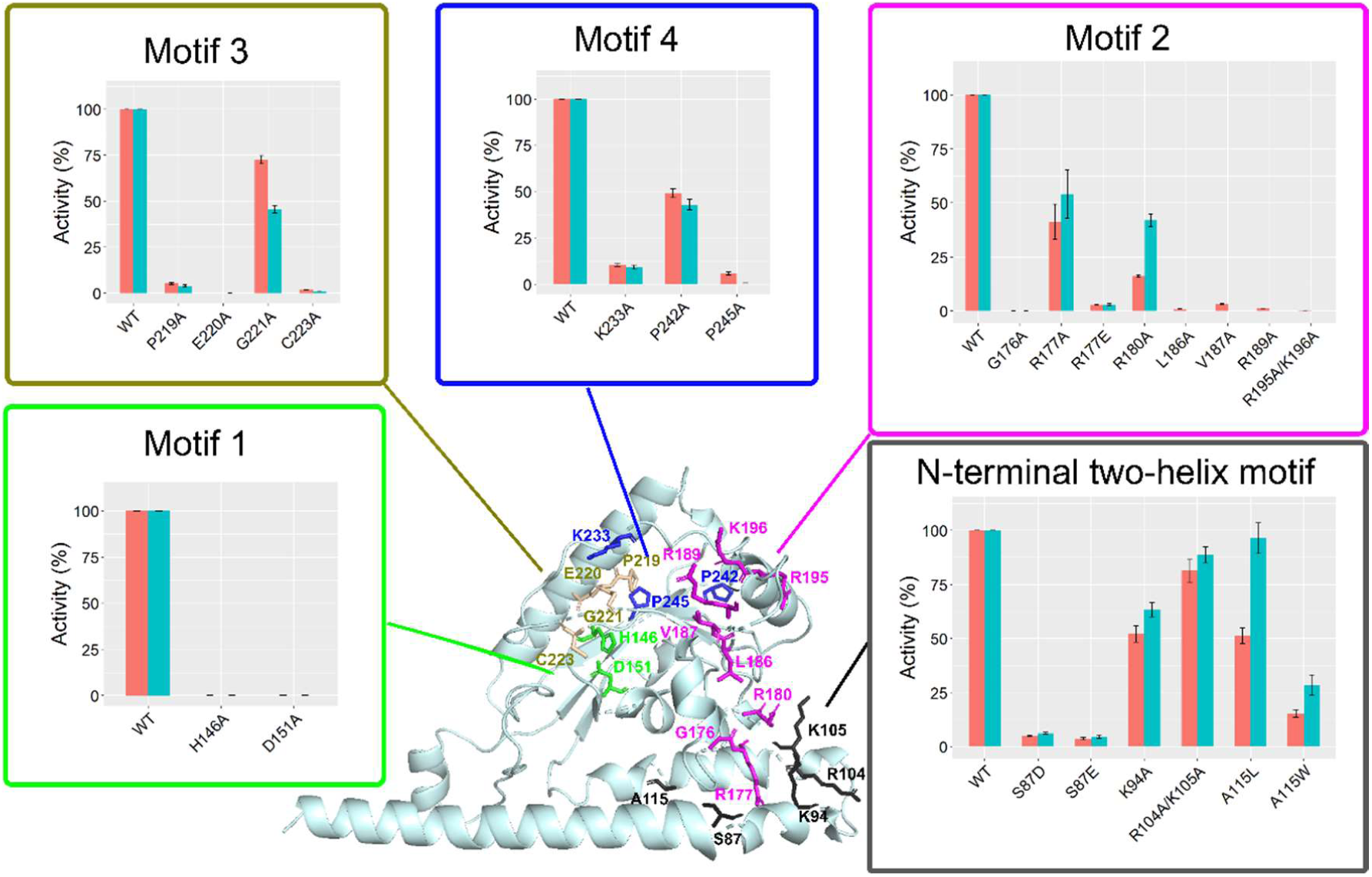
Analysis of conserved AGPAT motifs and the N-terminal two-helix motif. The positions of mutations tested experimentally are indicated in mLPCAT2 ribbon model, as follows: motif 1 mutants, green sticks; motif 2 mutants, magenta sticks; motif 3 mutants, blue sticks; motif 4 mutants, orange sticks; N-terminal two-helix motif mutants, black sticks. Enzyme activities using Acetyl-CoA (orange bars) or arachidonoyl-CoA (green bars) were measured for mLPCAT2 variants containing mutations of each motif. The results are illustrated as relative activities compared with the WT enzyme. The background activity was measured using vector-transfected cells and subtracted from the experimental values. The error bars represent the standard error (S.E.) of three independent experiments, each performed in triplicate.

**Fig. 3.**
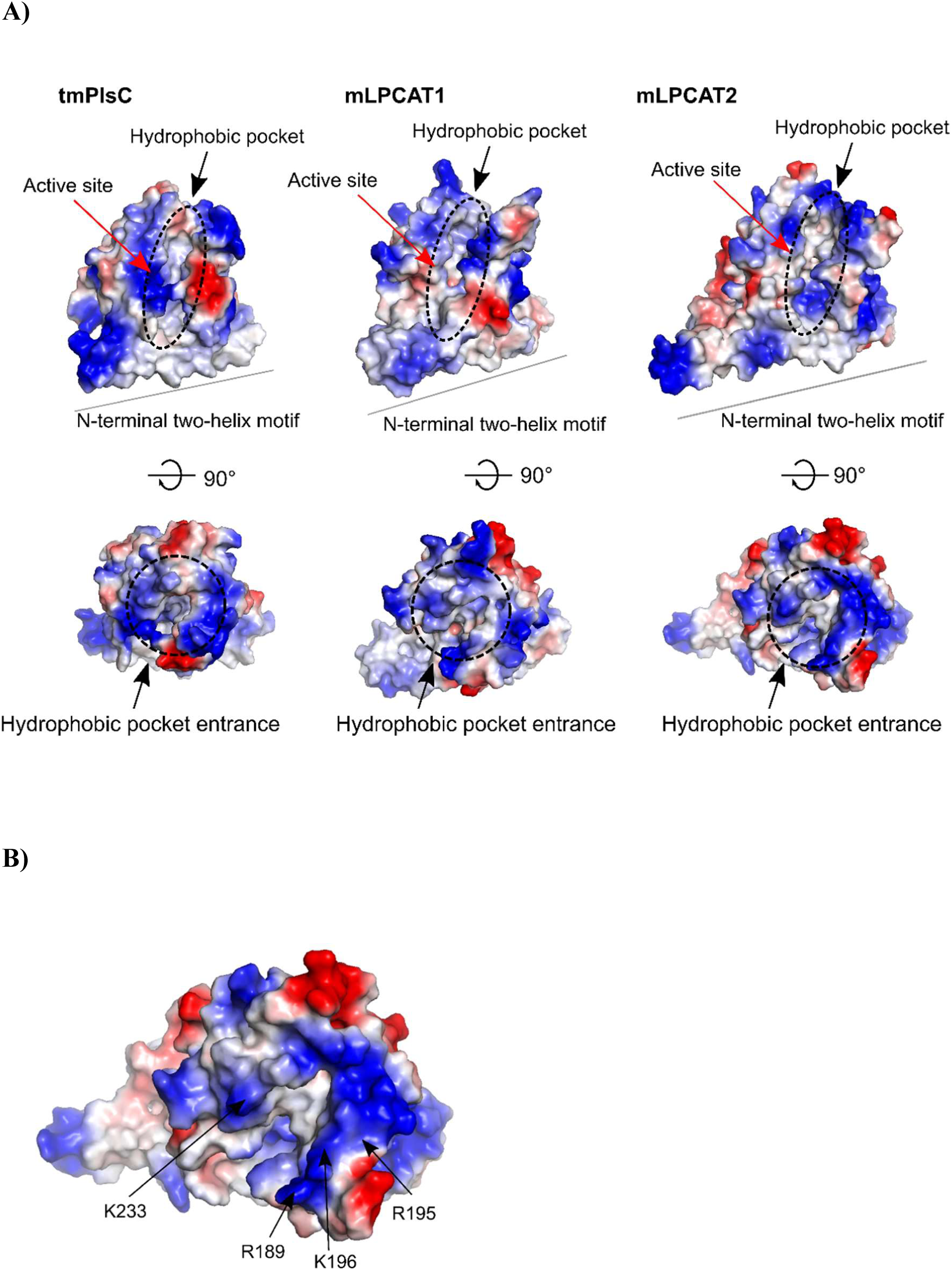
Electrostatic charges on the surfaces of tmPlsC, mLPCAT1, and mLPCAT2. A) Representations of the surface charges of tmPlsC, mLPCAT1, and mLPCAT2, showing the active site containing the H-X-X-X-X-D in motif 1, the hydrophobic pocket, the hydrophobic pocket entrance, and the N-terminal two-helix motif. B) Upper surface of mLPCAT2 on the cytoplasmic side. The positions of mutations studied around the entrance of the hydrophobic pocket are shown. The figures were drawn using PyMol software, version 2.3.2. Electrostatic charges were modeled using the default vacuum electrostatic package in PyMOL. In all electrostatic models, red shading represents negatively charged regions, blue shading represents positively charged regions, and white shading represents neutral/hydrophobic regions.

Previously, it was reported that the length of the hydrophobic pocket of tmPlsC determines its acyl-chain selectivity (18). It was also verified whether the size of the hydrophobic pocket of mLPCAT2 influences its acyl-chain selectivity. According to the model, Ser87 and Arg108 are on opposite sides of the N-terminal two-helix motif. Thus, Ser87 was substituted with Glu or Asp to examine whether the interaction between either amino acid would fill the space between the helices and affect the acyl-donor selectivity. Both the S87D and S87E mutants lost their lyso-PAF acyltransferase activities with acetyl-CoA and arachidonoyl-CoA (Fig. 2). S87E and S87D may bind to Arg108 and affect the structure of mLPCAT2. The predicted loop regions following motif 4 (Fig. 1A) were well conserved in mLPCAT1 and mLPCAT2. However, exchanging the loop of mLPCAT2 with that of mLPCAT1 resulted in the loss of lyso-PAF acetyltransferase and acyltransferase activities (Fig. 4).

**Fig. 4.**
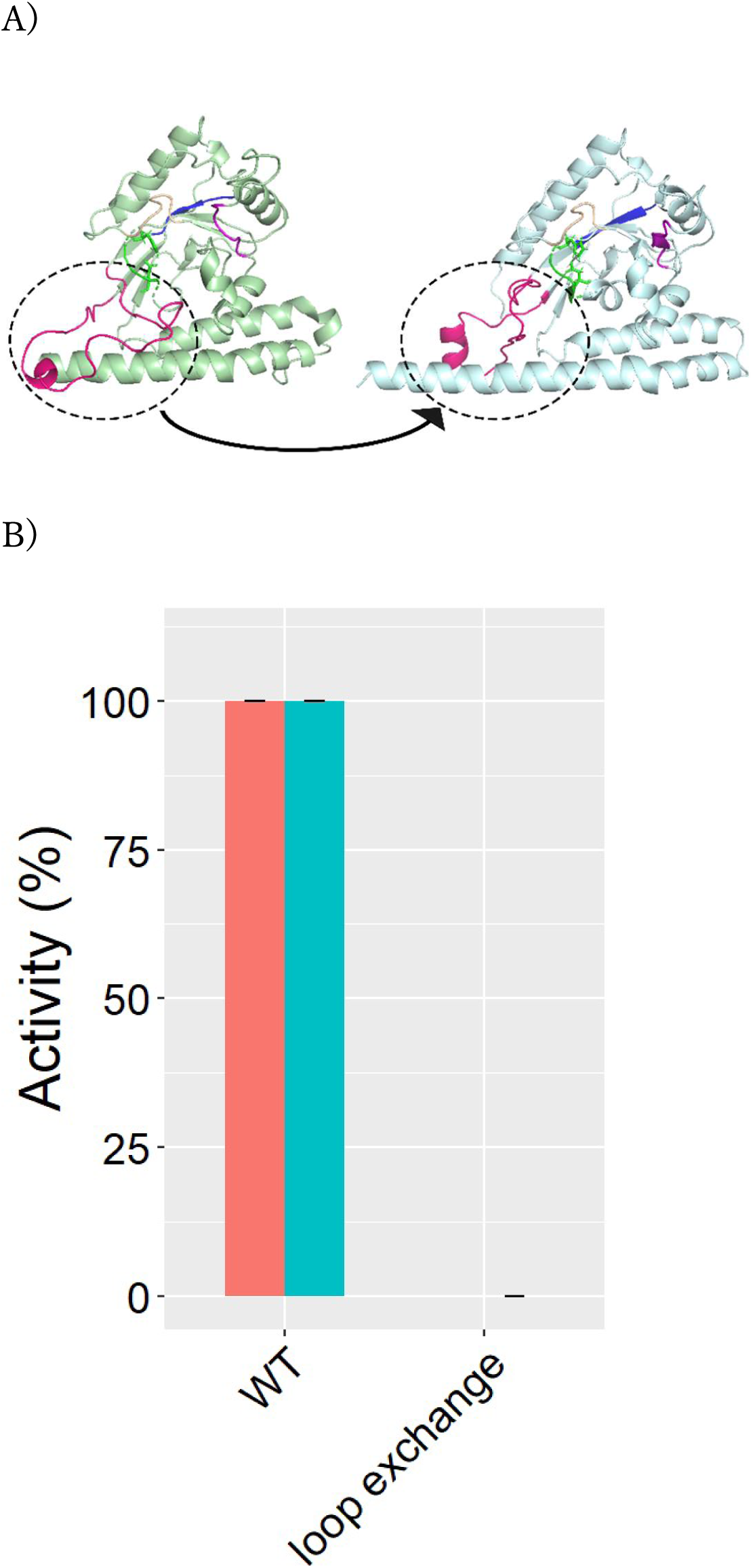
Analysis of the predicted loop region. A) Loop-exchange mutant with the loop sequence of mLPCAT2 exchanged for that of mLPCAT1. The loops are colored in magenta and are surrounded by a dashed line. The left ribbon model represents mLPCAT1, and the right model represents mLPCAT2. B) Enzyme activities using acetyl-CoA (orange bars) or arachidonoyl-CoA (green bars) were measured for the loop-exchange mutant of mLPCAT2.

### Acyl-donor selectivity

Next, Ala115 in the two-helix motif was substituted with Leu or Trp, which have bulky side chains (Fig. 5A). The A115L and A115W mutants remained partially active, but showed different preferences for the substrates, acetyl-CoA and arachidonoyl-CoA (Fig. 2). To evaluate acyl-donor selectivity, four different acyl-CoA mixtures, 16:0-, 18:1-, 18:2-, and 20:4-CoA, were used for the lyso-PAF acyltransferase assays (Fig. 5B). The substrate selectivity was also investigated for the R177A mutant, because Arg177 is thought to interact with the membrane (Fig. 5A). These protein-expression levels of each mutant were almost same (Supporting Fig. 1). The total amounts of alkyl-PCs (16:0/16:0, 16:0/18:1, 16:0/18:2, and 16:0/20:4) produced by each enzyme was set to 100% after subtracting the background signals of mock reactions. The acyl-donor substrate selectivity of each mutant was determined as the ratio of each product to the total quantity of alkyl-PC produced (Fig. 5B). The A115L and A115W mutants had lower preferences for 16:0-CoA and 20:4-CoA, respectively. In contrast, little change was observed in the acyl-donor preference of the R177A mutant. These results suggest that the amino acid size at this position may affect acyl-chain selectivity.

**Fig. 5.**
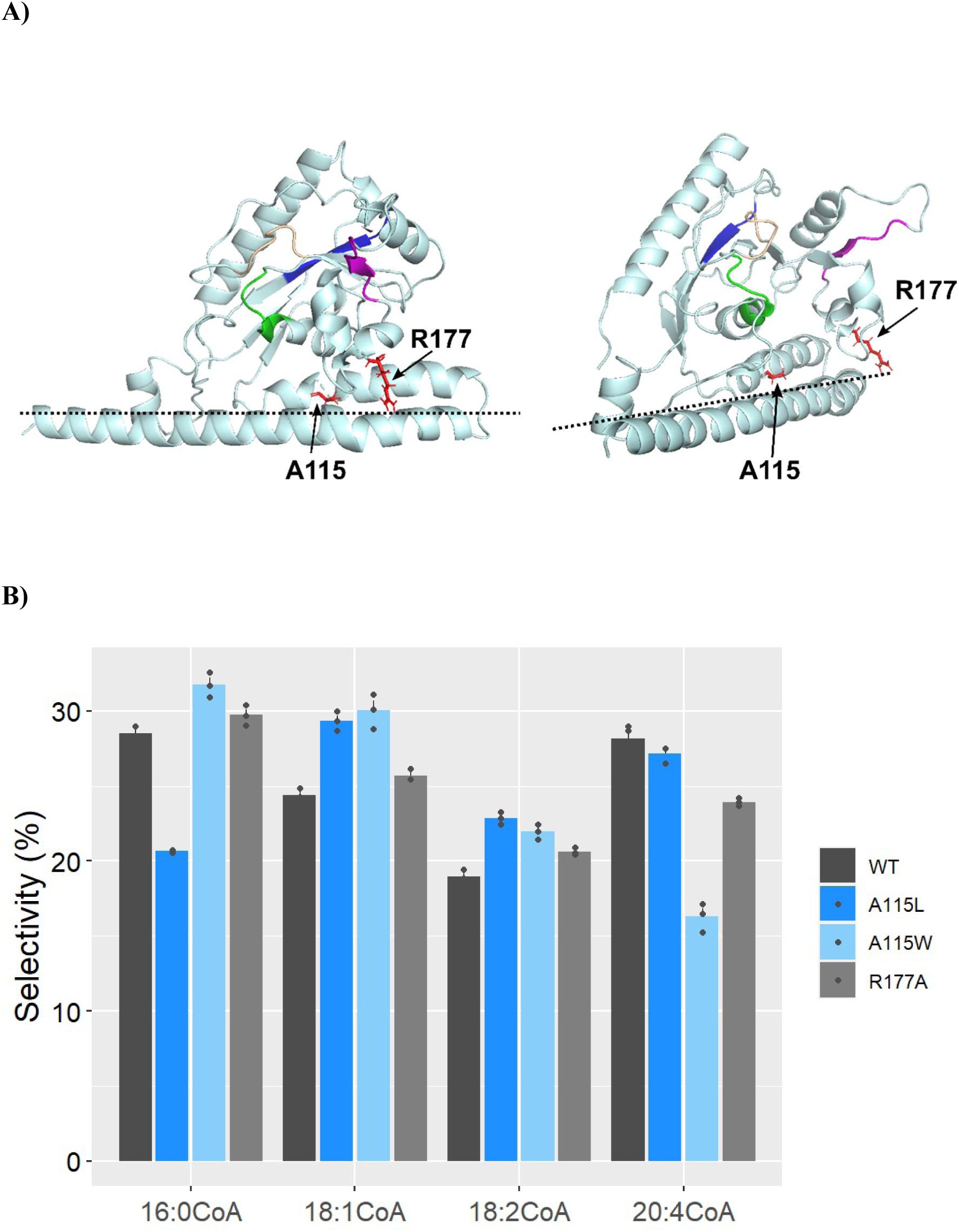
The effects of mutations in N-terminal two-helix motif on the acyl-donor selectivity. A) Location of A115 and R177 in mLPCAT2. A115 and R177 are represented with red sticks. The black dashed line indicates the predicted membrane location. B) The lyso-PAF acyltransferase activities of WT mLPCAT2 and A115L, A115W, and R177A mLPCAT2 mutants were measured using acyl-CoA mixture (16:0-CoA, 18:1-CoA, 18:2-CoA, and 20:4-CoA). Acyl-donor substrate selectivity was determined as the ratio of each product to the total amount of alkyl-PC. The error bars represent the S.E. of three independent experiments, each performed in triplicate.

### Acyl-CoA-binding model

The modeling results indicate that a hydrophobic pocket containing the active center, H-X-X-X-X-D (Motif 1) is positioned in the center of the enzyme. In addition, the AGPAT motifs, which have significant effects on the LPCAT2 activity detected by the mutant analysis, could exist around the pocket. Furthermore, Ala115, an amino acid that influences acyl chain selectivity, is predicted to be present in the bottom of the pocket. Thus, based on the homology-modeling results and mutant activities, acyl-CoA could fit into the hydrophobic pocket of mLPCAT2, similar to the PlsC model. This pocket is surrounded by four motifs, and the thioester moiety of the acyl-CoA fits near the active center (Fig. 6, Supporting Information model 2).

**Fig. 6.**
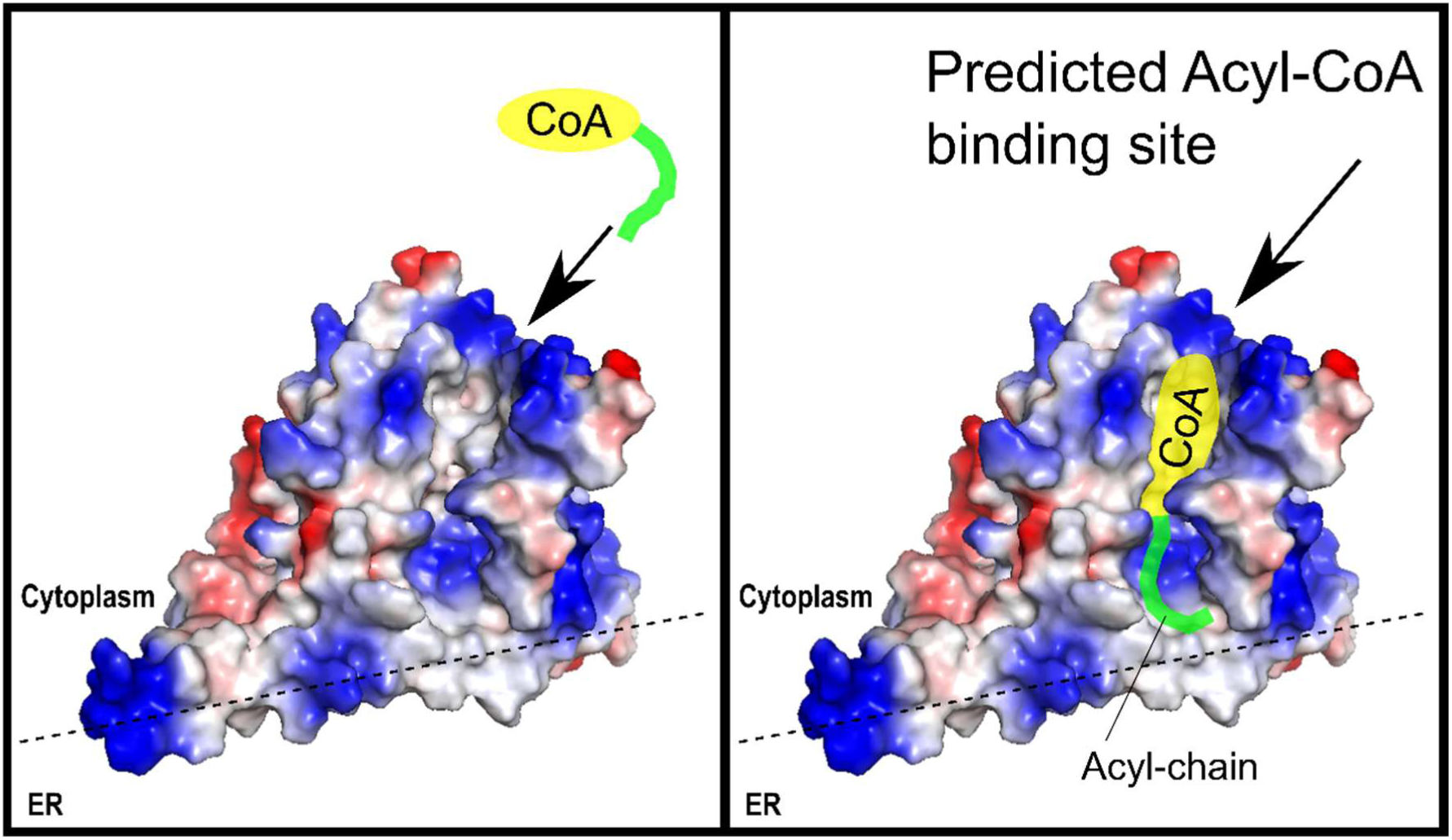
Acyl-CoA-binding model of mLPCAT2. The acyl-chain of acyl-CoA is colored in green, and CoA is colored in yellow. The black dashed line indicates the predicted membrane location. Acyl-CoA is thought to be recruited from the cytoplasm. The acyl-CoA-binding site is predicted to be located in the hydrophobic pocket, adjacent to positively charged residues at the top of the enzyme on the cytoplasmic side.

## Discussion

The data generated in this study provide the predicted structures of mLPCAT2 and mLPCAT1, based on homology modeling with tmPlsC as a reference. Our results showed that mLPCAT1 and mLPCAT2 have two N-terminal α-helices, which is known to contact the cellular membrane. Mutagenesis showed that the four AGPAT motifs of mLPCATs were essential for mLPCAT2 activities and substrate preferences. We identified a potential binding pocket for acyl-CoA. The four AGPAT motifs of mLPCAT2 were located around the binding pocket. Our data highlight the mLPCAT2 structure and may contribute to the development of LPCAT2 inhibitors.

Based on sequences registered in the PDB, several enzyme structures have been predicted to be monotopic membrane proteins, such as cytochrome c oxidase subunit II (20) and carnitine palmitoyltransferase 2 (21). These enzymes interact with amphipathic substrates within the phospholipid bilayer or in the cytosol (22). Our results are consistent with a previous report indicating that mLPCAT1 and mLPCAT2 are monotopic membrane proteins (13). The donor substrates of mLPCAT2 are the amphipathic molecule, acyl-CoA, or the hydrophilic molecule, acetyl-CoA. The acceptor molecule, lyso-PAF (LPC), is present in the membrane. The monotopic structure of mLPCAT2 might be beneficial for using two distinct substrates.

Our homology-modeling data showed that motifs 1–4, which are highly conserved in the AGPAT family, appeared to surround a hydrophobic pocket around the active center. Several positively charged amino acids were localized on the protein surface around the entrance of the hydrophobic pocket, on the cytoplasmic side and near the endoplasmic lipid membrane. Mutagenesis revealed that mutations near the active center were likely to lead to the loss of the catalytic activity and that substituting Ala for positively charged amino acids near the hydrophobic pocket or the membrane reduced the activities of mLPCAT2. Moreover, because the A115L and A115W mutations changed the acyl-chain substrate selectivity, Ala115 may be important for discriminating between different acyl-CoA molecules. Further studies, such as calculating *K*m values, are needed to compare the substrate preferences of mLPCAT2. In addition to affecting the active center, point mutations reduced the enzymatic activity of mLPCAT2, potentially by changing the interactions of the enzyme with substrates and/or affecting the stability of the enzyme structure.

The effects of point mutations on mLPCAT1 activities have already been reported (6). mLPCAT1 also possesses the four AGPAT motifs. By performing homology modeling of mLPCAT1 with tmPlsC, we found that the predicted acyl-CoA-binding pocket of mLPCAT1 was surrounded by the AGPAT motifs (Fig. 1C, Supporting Information model 1). Highly conserved motifs in the AGPAT family are critical regions for acyl-CoA binding. Further studies are needed to identify the regions in mLPCAT1 and mLPCAT2 that react with lyso-PAF. Because lyso-PAF is located within membrane, the N-terminal two-helix motif may play important roles in the reaction with lyso-PAF.

In this study, homology modeling with the crystal structure of tmPlsC revealed a potential acyl-CoA-binding pocket in mLPCAT2. This area was surrounded by the AGPAT motifs and the acyl chain may be coming from the cytosol. Although this is a first report to present an analysis of the mLPCAT2 homology model structure and it is important to understand biochemical characteristics of mLPCAT2, it is necessary to further examine the mLPCAT2 crystal structure in the future. We studied shortened structures of mLPCAT2 and mLPCAT1 because the full-length tmPlsC sequence is shorter than those of them. Thus, Ser34 at the N-terminus of mLPCAT2 was not analyzed in the data presented in Fig. 1A and B. Because mLPCAT2 is phosphorylated and activated in response to extracellular stimuli, determining the full-length structure of mLPCAT2 (including Ser34) is important for understanding the PAF-biosynthesis mechanism under inflammatory conditions. Further studies are also needed on this topic.

In summary, we determined homology modeling-based structures for the acyl-CoA-binding pockets of mLPCAT1 and mLPCAT2. Our findings indicate that four AGPAT motifs surround this pocket (Fig. 1C, Supporting Information models 1, 2). Thus, the fatty acid of acyl-CoA, which is a donor for mLPCAT2, was localized near the active center. Acyl-CoA is thought to be recruited from the cytosol (Fig. 6). In contrast, mLPCAT2 recruits lyso-PAF (an acceptor for mLPCAT2) from the endoplasmic reticulum membrane, as mLPCAT2 is a monotopic protein. Studying and understanding the structure of mLPCAT2 paves the way for analyzing the biosynthetic mechanism of PAF and for developing medicines to overcome PAF-related conditions, such as neuropathic pain, inflammation, and allergy.

## Experimental Procedures

### Materials

Lyso-PAF C16-d4 and d4-PAF were purchased from Cayman Chemical Co. (Ann Arbor, MI, USA). Dimyristoyl-PC (DMPC), LPC, alkyl 36:4 PC, 16:0-CoA, 18:1-CoA, 18:2-CoA, and 20:4-CoA were obtained from Avanti Polar Lipids (Alabaster, AL, USA). 2:0-CoA, liquid chromatography–mass spectrometry (LC–MS)-grade acetonitrile, high-performance liquid chromatography-grade methanol, isopropanol, chloroform, and ammonium bicarbonate were obtained from FUJIFILM Wako Pure Chemical Corp. (Osaka, Japan). Protein Assay Dye Reagent Concentrate was obtained from Bio-Rad (Hercules, CA, USA). Bovine serum albumin (BSA) was purchased from Sigma Aldrich (St. Louis, MO, USA).

### Mutagenesis of mLPCAT2

Each mLPCAT2 mutant was produced by overlap-extension PCR (23) using KOD-Plus-(TOYOBO; Osaka, Japan). The sequences of the primer sets used for overlap-extension PCR are shown in Table S2. Amplified PCR products were cloned into the pCXN2.1 vector and sequenced.

### Preparation of microsomal proteins from Chinese hamster ovary-K1 (CHO-K1) cells

CHO-K1 cells in 10-cm dishes were transfected with complementary DNA encoding FLAG-tagged mLPCAT2 or one of several FLAG-tagged mLPCAT2 mutants using Lipofectamine 2000 (Thermo Fisher Scientific, USA). At 48 h post-transfection, the cells were scraped into 1 mL of ice-cold buffer containing 20 mM Tris-Cl (pH 7.4), 300 mM sucrose, and 1× cOmplete protease inhibitor cocktail (Sigma Aldrich), and then sonicated three times on ice, for 30 s each time. After certification at 9,000 × *g* for 10 min, each supernatant was collected and centrifuged again at 100,000 × *g* for 1 h. The resulting pellets were resuspended in 300 µL of buffer containing 20 mM Tris-Cl (pH 7.4) and 1 mM ethylenediaminetetraacetic acid (EDTA). Protein concentrations were determined using the Bradford method, using Protein Assay Dye Reagent Concentrate (Bio-Rad) and BSA (essentially fatty acid-free lyophilized powder; Sigma) as a standard.

### Measurement of lyso-PAF acetyltransferase (lyso-PAF-2:0-CoA assay) and lyso-PAF acyltransferase (lyso-PAF-20:4-CoA assay) activities

Microsomal protein (0.05 µg) was added to reaction mixtures containing 100 mM Tris-Cl (pH 7.4), 1 mM EDTA, 1 mM CaCl2, 0.015% Tween-20, and 5 µM lyso-PAF C16-d4 as an acceptor, and 1 mM 2:0-CoA (for lyso-PAF-2:0-AT assays) or 25 µM 20:4-CoA (for lyso-PAF-20:4-AT assays) as an acyl-donor, in a total volume of 0.1 mL. After incubation at 37°C for 10 min, the reactions were stopped by the addition of 0.3 mL of methanol. As an internal standard, 10 µL of 1 µM 17:0 LPC or 5 µM DMPC was added for the lyso-PAF-2:0-AT or lyso-PAF-20:4-AT assays, respectively. After centrifugation at 10,000 × *g* for 5 min, the supernatants were analyzed by LC-MS/MS (Waters, Milford, MA and Thermo Scientific, Waltham, MA) to quantify each product (d4-PAF or d4-alkyl 36:4 PC).

### Measuring the activities of lyso-PAF acyltransferase in lyso-PAF-acyl-CoA-mixture assays

Microsomal protein (0.05 µg) was added to reaction mixtures containing 100 mM Tris-Cl (pH 7.4), 1 mM EDTA, 1 mM CaCl2, 0.015% Tween-20, and 5 µM lyso-PAF C16-d4 as an acceptor, and 5 µM each of 16:0-, 18:1-, 18:2- and 20:4-CoA (all from Avanti) as an acyl-donor, in total volume of 0.1 mL. After incubation at 37°C for 10 min, the reactions were stopped by adding 0.3 mL of chloroform/methanol (MeOH) (1:2, v/v). Lipid extraction was performed using the Bligh and Dyer method, after which 50 µL of 0.02 µM DMPC in MeOH was added as an internal standard. Each lipid extract was dried and dissolved in 0.1 mL of MeOH and analyzed by LC-MS/MS (LCMS8050, Shimadzu Corp. Kyoto, Japan) to measure the peak areas of d4-alkyl PC.

### LC-MS/MS analysis of phospholipids

Phospholipids (products of the lyso-PAF-2:0-CoA or 20:4-CoA assays) were analyzed using an LC-MS/MS system composed of an Acquity UPLC system (Waters, Milford, MA, USA) and a triple quadrupole TSQ Vantage MS instrument (Thermo Scientific) (24). A BEH C8 column (2.1 × 30 mm; Waters) was used for the chromatographic separation of PAF and PC. Solvent A1 was 20 mM ammonium bicarbonate (Wako) and solvent B1 was acetonitrile (Wako). For PAF, the pump gradient [time(%A/%B)] was programmed as follows: 0 min (80/20)-4.5 min (5/95)-6.0 min (5/95)-6.01 min (80/20). For PC, the pump gradient [time(%A/%B)] was programmed as follows: 0 min (80/20)-3.0 min (30/70)-3.01 min (5/95)-4.25 min (5/95) −4.26 min (80/20). The flow rate was set to 800 µL/min, and 5 µL of each sample was injected.

Phospholipids (products of the lyso-PAF-acyl-CoA-mixture assays) were analyzed using an LC-MS/MS system composed of a Nexera Ultra High-Performance Liquid Chromatograph system (Shimadzu Corp., Kyoto, Japan) and a triple quadrupole mass spectrometer (LCMS-8050, Shimadzu). An Acquity UPLC BEH C8 (2.1 × 100 mm; Waters) column was used for reversed-phase liquid chromatography with three phases: 5 mM ammonium bicarbonate (solvent A), acetonitrile (solvent B), and isopropanol (solvent C). The pump gradient [time(%A/%B/%C)] was programmed as follows: 0 min (70/15/15)-1.5min (20/40/40)-8 min (20/40/40)-8.1 min (6/47/47)-15.0 min (6/47/47)-15.1min (70/15/15)-18:00 (70/15/15). The flow rate was set to 300 µL/min, and 5 µL of each sample was injected. Selected reaction monitoring (SRM) transitions used for the analysis are provided in a Table S1.

### Western blot analysis

Microsomal protein (2 µg) was separated by 10% sodium dodecyl sulfate-polyacrylamide gel electrophoresis and transferred to nitrocellulose blotting membranes (GE Healthcare Life Sciences, Little Chalfont, UK). The membranes were blocked with 5% skim milk (BD Biosciences, San Jose, CA, USA) in Tris-based buffer containing 0.1% Tween-20 (Wako, Osaka, Japan), incubated with mouse M2 anti-FLAG monoclonal antibody (1:1000) (Sigma), washed with Tris-based buffer containing 0.1% Tween-20, and then incubated with horseradish peroxidase-linked anti-mouse IgG (1:1000) (GE Healthcare Life Sciences). After washing, the membranes were exposed to the ECL Select Western Blotting Detection Reagent (GE Healthcare Life Sciences). The reactive proteins were detected using Image Quant LAS500 software (GE Healthcare Life Sciences).

### Homology modeling

We performed LPCAT homology-model calculations using the crystal structure of tmPlsC (PDB code 5KYM) and the MODELLER program (25) in Discovery Studio 2018. Predictions generated using the PSIPRED algorithm (26) indicated that similar secondary structures existed between tmPlsC, mLPCAT1, and mLPCAT2. Sequence identities and similarities were confirmed using ClustalW software (27). The constructed model was selected based on the probability density functions total energy score and Discrete Optimized Potential Energy analysis (28). Furthermore, the geometry of the model was optimized using CHARMM (29) and NAMD (30) in Discovery Studio with harmonic restraint, using the default settings.

### Software

Sequence alignments were generated using GENETYX-MAC software version 17.0.2 (GENETYX Corporation). For the alignments, the sequences of mLPCAT1 (NP_663351), mLPCAT2 (NP_766602), and tmPlsC (WP_004082209.1) were obtained from the National Center for Biotechnology Information database. All statistical calculations were performed using the R statistical package, version 4.0.0 (31).

## Acknowledgements

We are grateful for constructive comments from K. Waku (Teikyo University). We also thank Dr. J.-I. Miyazaki (Osaka University) for supplying the pCXN2.1 vector. We thank M. Yamamoto and Y. Takahashi (National Center for Global Health) for their excellent technical assistance. We would like to thank all members of the Shimizu laboratories (National Center for Global Health and Medicine and the University of Tokyo) for their advice, discussion, and technical support.

## Author contributions

Fumie Hamano designed the study, performed the experiments, analyzed the data, and wrote the manuscript. Kazuaki Matoba and Nobuo N. Noda performed the homology modeling. Tomomi Hashidate-Yoshida and Tomoyuki Suzuki performed mLPCAT2 assays. Takeshi Harayama designed the study and analyzed the data. Kiyotake Miura generated the plasmids encoding mLPCAT2 mutants. Daisuke Hishikawa and Koichi Yuki supported the experiments and interpreted the results. Yoshihiro Kita analyzed the data. Hideo Shindou and Takao Shimizu designed and wrote the manuscript. All authors assisted in editing the manuscript.

## Funding and additional information

This work was supported by AMED-CREST (grant number JP20gm0910011 to HS), AMED-P-CREATE (grant number JP20cm0106116 to HS), and the AMED Program for Basic and Clinical Research on Hepatitis (grant number JP20fk0210041 to HS). TS was supported by the Takeda Science Foundation. TH was supported by the French Government (National Research Agency, ANR) through the “Investments for the Future” programs LABEX SIGNALIFE ANR-11-LABX-0028 and IDEX UCAJedi ANR-15-IDEX-01, and the ATIP-Avenir program (CNRS/Inserm).

The current academic positions of Kiyotake Miura is Shuyukan senior high school.

## Abbreviations

AGPAT: 1-acylglycerol-3-phosphate O-acyltransferase
CoA: coenzyme A
DMPC: dimyristoyl-PC
EDTA: ethylenediaminetetraacetic acid
LC: liquid chromatography
LPCAT: lysophosphatidylcholine acyltransferase
MeOH: methanol
mLPCAT: mouse LPCAT
MS: mass spectrometry
PAF: platelet-activating factor
PC: phosphatidylcholine
PDB: Protein Data Bank
S.E.: standard error
SRM: selected reaction monitoring
WT: wild-type

---

**Table S1:**
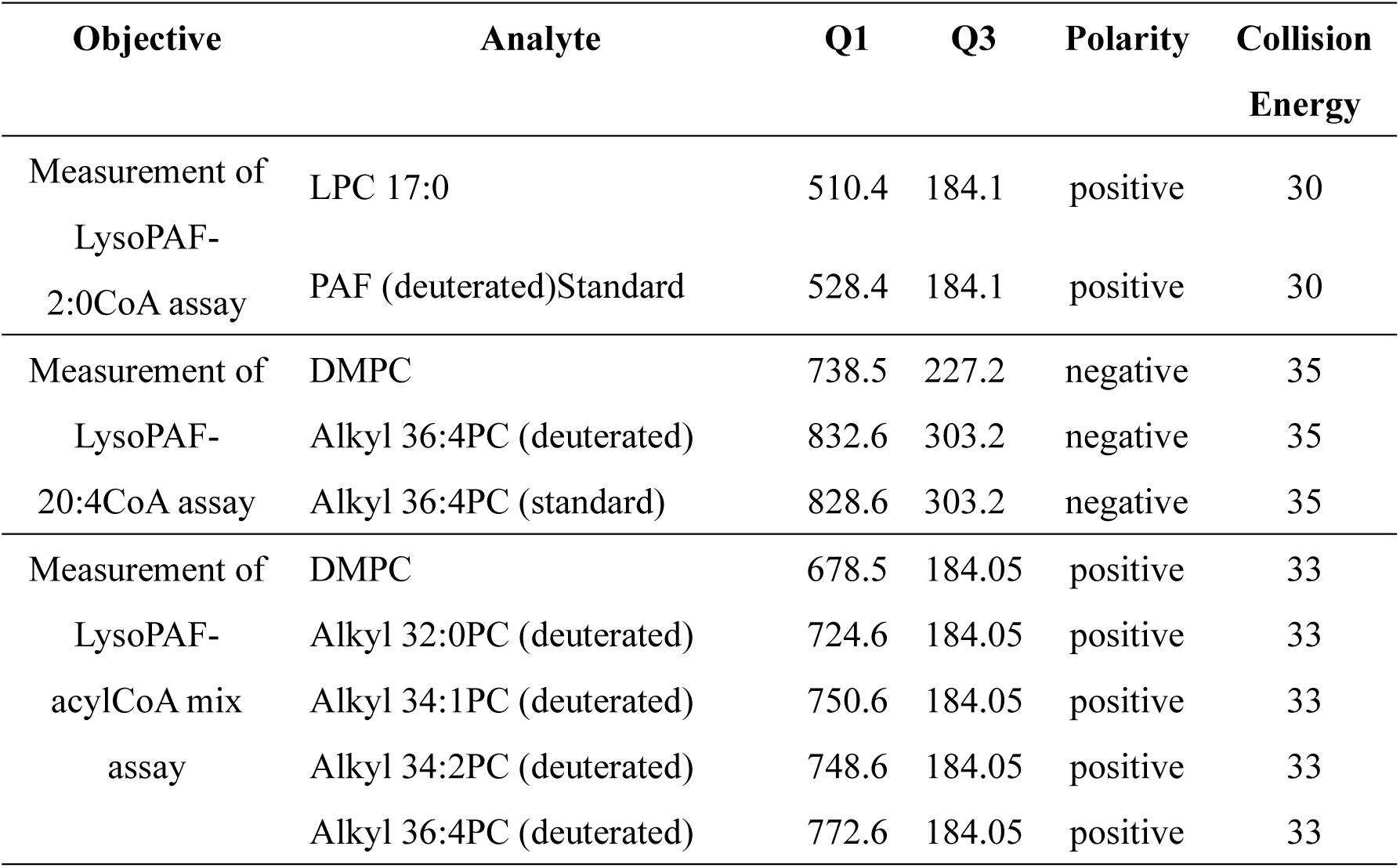
SRM transition used for LC-MS/MS analysis.

**Table S2:**
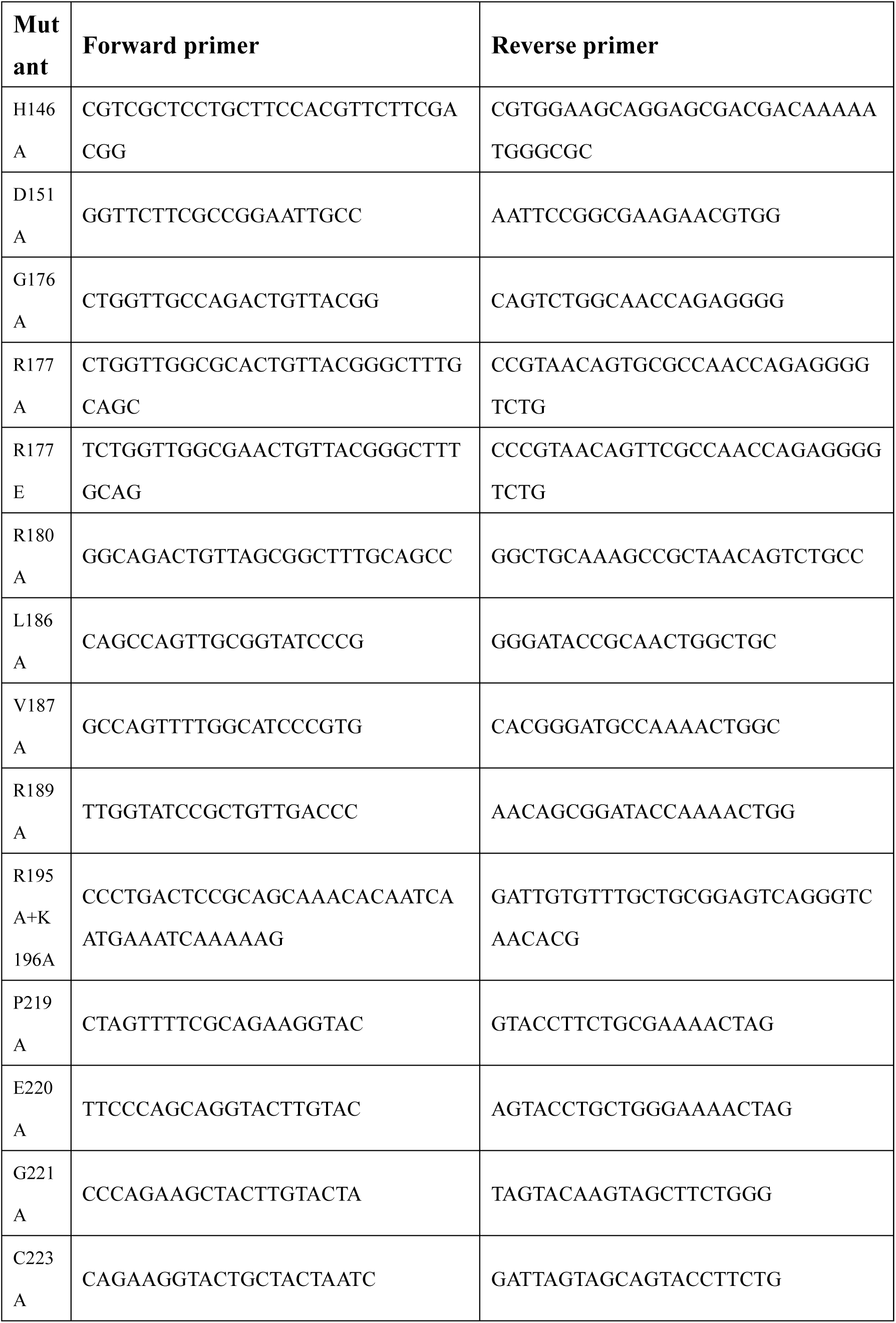
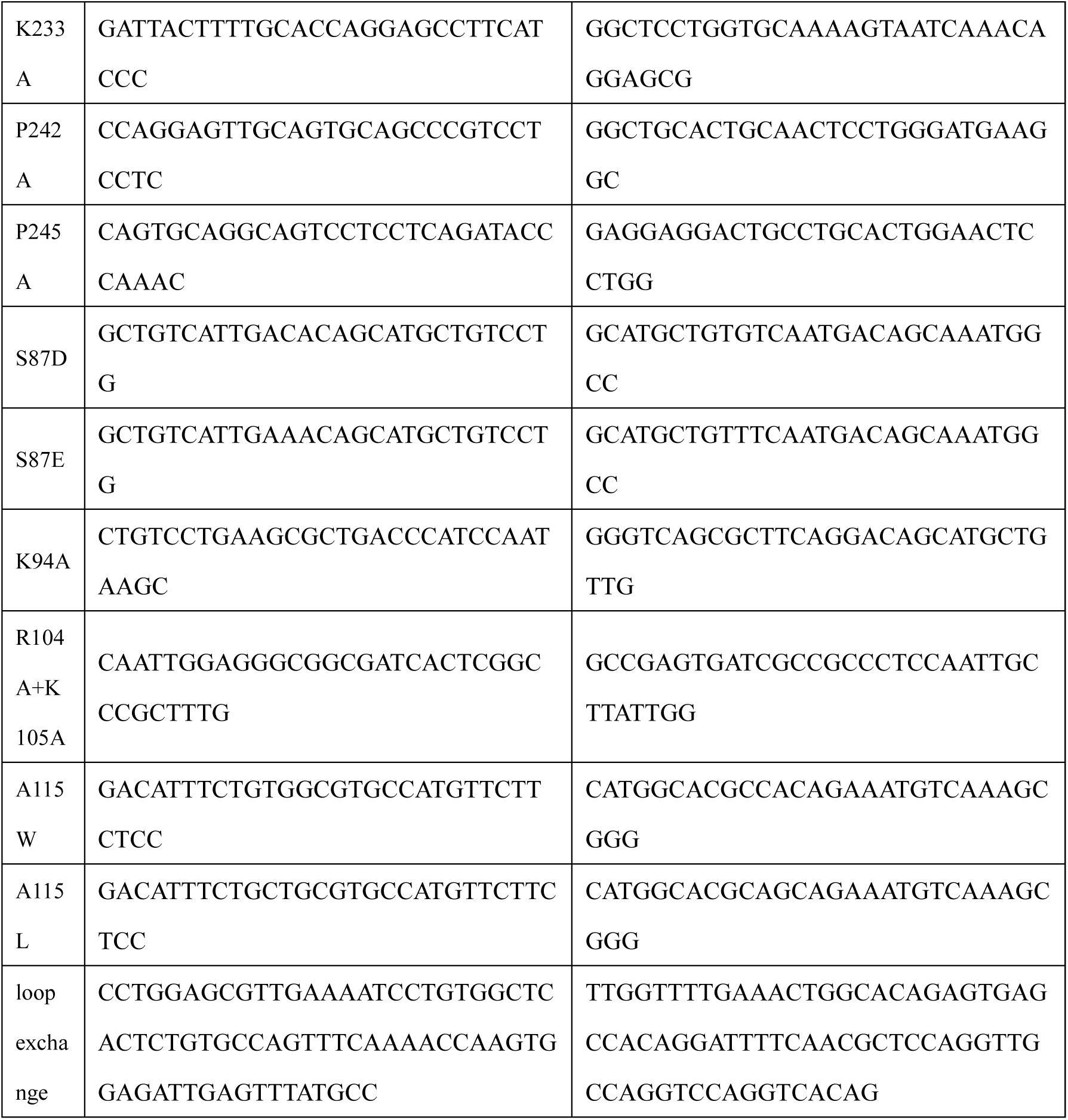
Primer sequences for generating mLPCAT2 mutants.

**Supporting Fig. 1.**
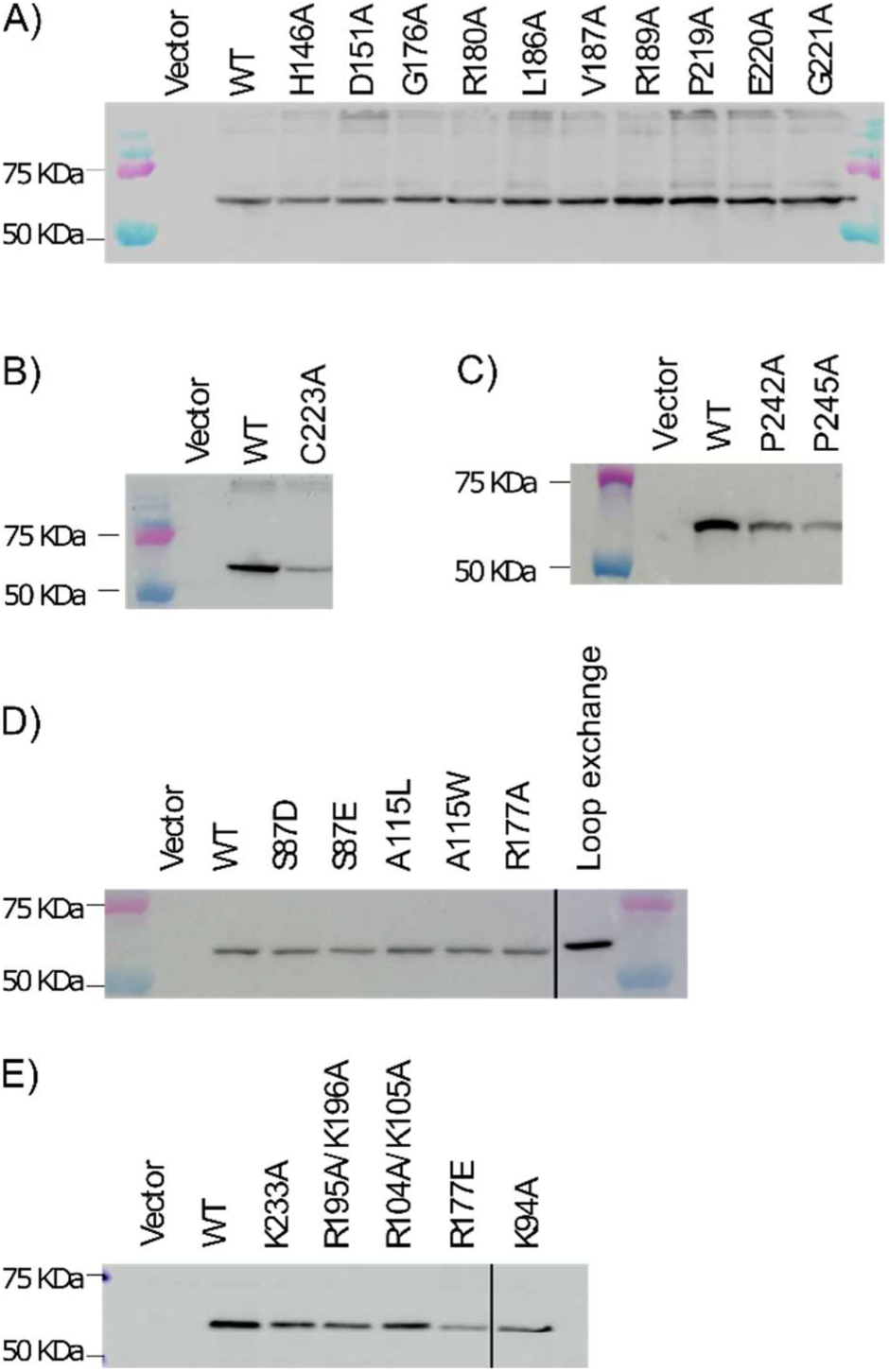
Confirmation of the expression of mLPCAT2 mutants by western blotting using an anti-FLAG antibody. Western blots were performed as described in the Experimental Procedures section. Vector, empty vector expression; WT, wild-type mLPCAT2. No signal was detected in cells transfected with the empty vector, whereas bands of expected size were observed in cells expressing WT mLPCAT2 or mLPCAT2 mutants. Three independent experiments were performed with similar results. These photos were cut to remove unused proteins, which are indicated by black lines.

**Supporting Fig. 2.**
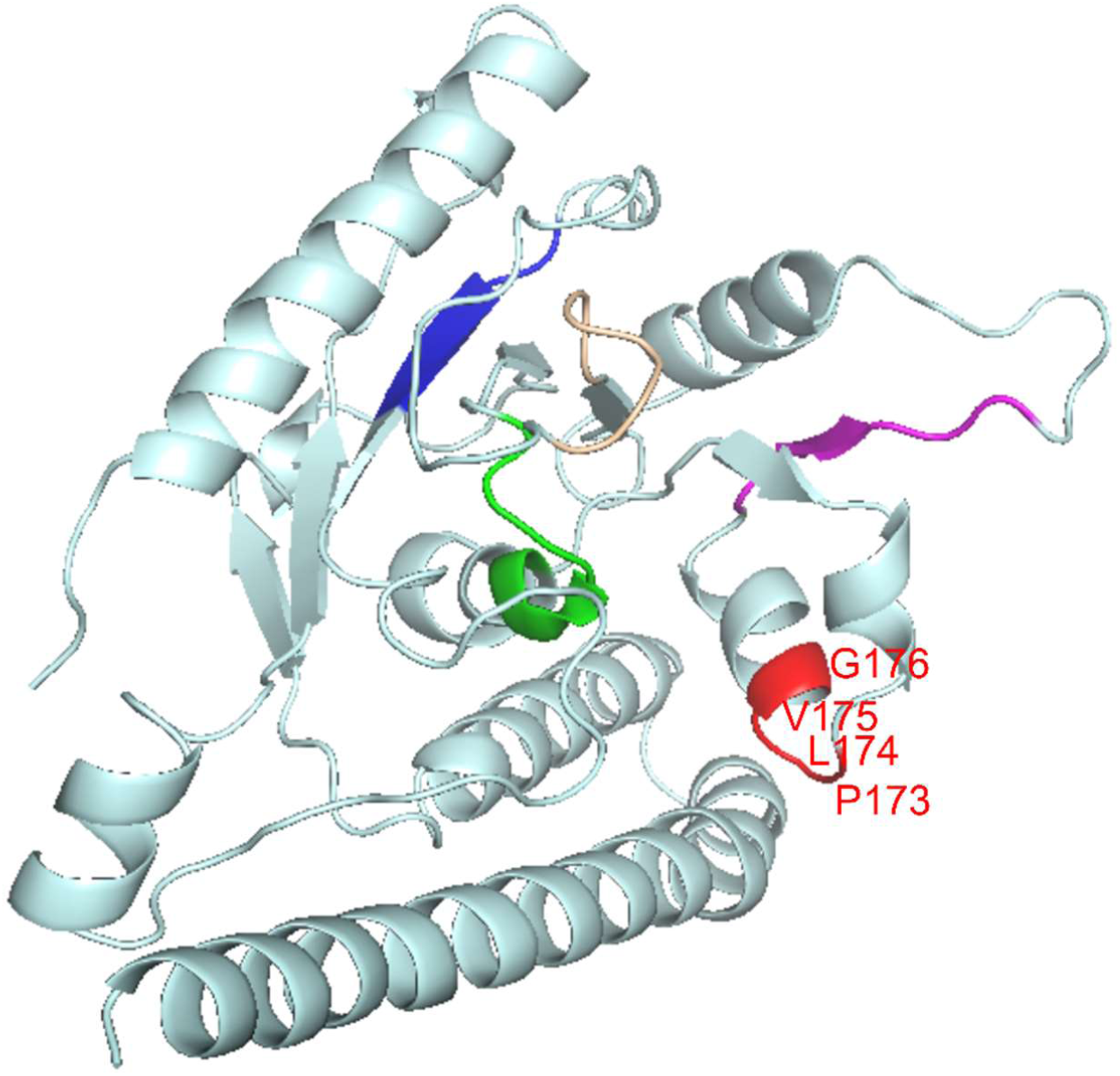
Location of the P-X-X-G sequence in the mLPCAT2 model. The Pro173-x174-x175-Gly176 is highly conserved among various LPLATs and could introduce a kinked structure between motif 1 and motif 2. The P-X-X-G sequence is colored in red.

